# Organization of the Proteostasis Network of Membraneless Organelles

**DOI:** 10.1101/2024.11.12.623332

**Authors:** Christine M. Lim, Yuqi Bian, Alicia González Díaz, Frank Pun, Alex Zhavoronkov, Richard I. Morimoto, Michele Vendruscolo

## Abstract

Membraneless organelles (MLOs) are dynamic macromolecular condensates that act as crucibles to modulate biological process within the cell. Since MLOs form in the absence of lipid membranes, it is important to understand how their effective regulation is achieved by the protein homeostasis (proteostasis) system. To address this question, we report a comprehensive mapping of the proteostasis network (PN) of MLOs, comprising over 220,000 protein-protein interactions. The analysis of this PN reveals how the regulatory proteins (PN proteins) occupy central roles in the overall protein-protein interaction network of MLOs. We then investigate which branches of the PN are most important in the regulation of MLOs, and find that the anabolic component, which makes up ∼30% of the PN, is more closely involved with MLOs than the catabolic component, which makes up the remaining ∼70% of the PN. We also find that translation-related PN proteins and molecular chaperones play central roles in the regulation of MLOs. We then analyse the similarity between the PNs of different MLOs, revealing that stress granules and P-bodies share large portions of their PNs. Finally, to understand how specificity is achieved despite these similarities, we further analyse the PN organization, showing that HSP70 chaperones are generic in regulating MLOs, while client-specific HSP70 co-chaperones confer specificity to the chaperone action. Taken together, these results identify the composition of the PN of MLOs, rationalize its organization, and reveal a central role of molecular chaperones in the regulation of protein conformations within MLOs.

## Introduction

Membraneless organelles (MLOs) are macromolecular assemblies that lack lipid membranes but are crucial for orchestrating key cellular processes such as gene regulation, RNA metabolism and response to stress^1-5^. Given their central roles in managing cellular homeostasis, the improper assembly and disassembly of MLOs may impair the optimal functioning of the cell, contributing to pathological phenomena^4-7^. For example, the dysregulated assembly of stress granules has been implicated in cancer, viral infection, and neurodegeneration, while the aberrant formation of nucleoli was found to occur in ageing and in diverse disease conditions^4-7^. Hence, the regulation of the macromolecular components of MLOs is essential for optimal cellular function.

In this context, the protein homeostasis (proteostasis) network (PN)^8,9^ - the ensemble of proteins and protein-protein interactions involved in biological processes responsible for the expression and maintenance of the functional proteome^10,11^ - is key in regulating the formation, clearance, composition, localization and physical properties of MLOs^12-16^. Since MLOs are not bound by lipid membranes, and yet their mechanism of self-assembly appears to be robust, one would like to understand how the different MLOs are specifically regulated to ensure that they reliably assemble and disassemble as they need to.

In an early study of the mechanisms of regulation of MLOs, morphological changes of nucleoli, Cajal bodies, splicing speckles, PML nuclear bodies, cytoplasmic processing bodies, and stress granules were monitored upon down-regulation of 1354 genes^17^. Based on that analysis, 80% of the targeted genes were found to modify the morphology of MLOs. However, approximately 70% of the genes identified in that study affect multiple MLOs (**Figure S1**), underscoring the complexity of the task of understanding the specific mechanisms at play in the regulation of individual MLOs by the PN.

To address this problem, we present a large-scale mapping of the PN of MLOs and study its organization. We show that proteins in the PN (PN proteins) occupy central positions in the protein-protein interaction (PPI) networks of MLOs. By using the taxonomic structure recently proposed for the PN^8,9^, we then investigate which functional branches of the PN are most important in the regulation of MLOs, finding that a large proportion of translation-related proteins and molecular chaperones are closely associated with MLOs. Next, we study the similarity between the PN regulation of different MLOs. A comparison of the PNs of different MLOs reveals that stress granules and P-bodies share large portions of their individual PNs. Finally, we find that HSP70 proteins^11,18^ are essential in modulating MLO disassembly, with HSP70 exerting promiscuous activity on MLO regulation while the co-chaperones of HSP70 enable specificity. This is in line with prior reports the role of HSP70 on MLO dynamics^19-23^. Overall, the mapping of the PN of MLOs that we report, together with its initial analysis, offer a tool for the investigation of the mechanism of regulation of these important cellular structures. All data used in this study were from comprehensive LLPS data resources.

## Results

### Mapping of the proteostasis network (PN) of membraneless organelles

The PN of MLOs described in this work is a list of pairwise interactions in which the first protein is a component of an MLO and the second protein has been reported to interact with the first. To generate a comprehensive mapping of the PN of MLOs, we started from the protein-protein interaction networks of 6 well-characterized MLOs (stress granules, P-bodies, nucleoli, PML nuclear bodies, centrosomes, and post-synaptic densities (PSDs)) (**Supplementary Data 1)**. For this purpose, we employed human protein-protein interaction data from BioGRID^24^. Then, known interactions between proteins localising in each MLO were obtained from DrLLPS^25^, and used to create MLO-specific PNs (**Figure 1A**). Examples of PNs of individual MLOs are shown in **Figure S2A-D**. We then created an overall interaction network by pulling together all the MLO-specific PNs. Using this approach, we found that many PN proteins are shared among multiple MLOs (**Figure 1B**).

**Figure 1.**
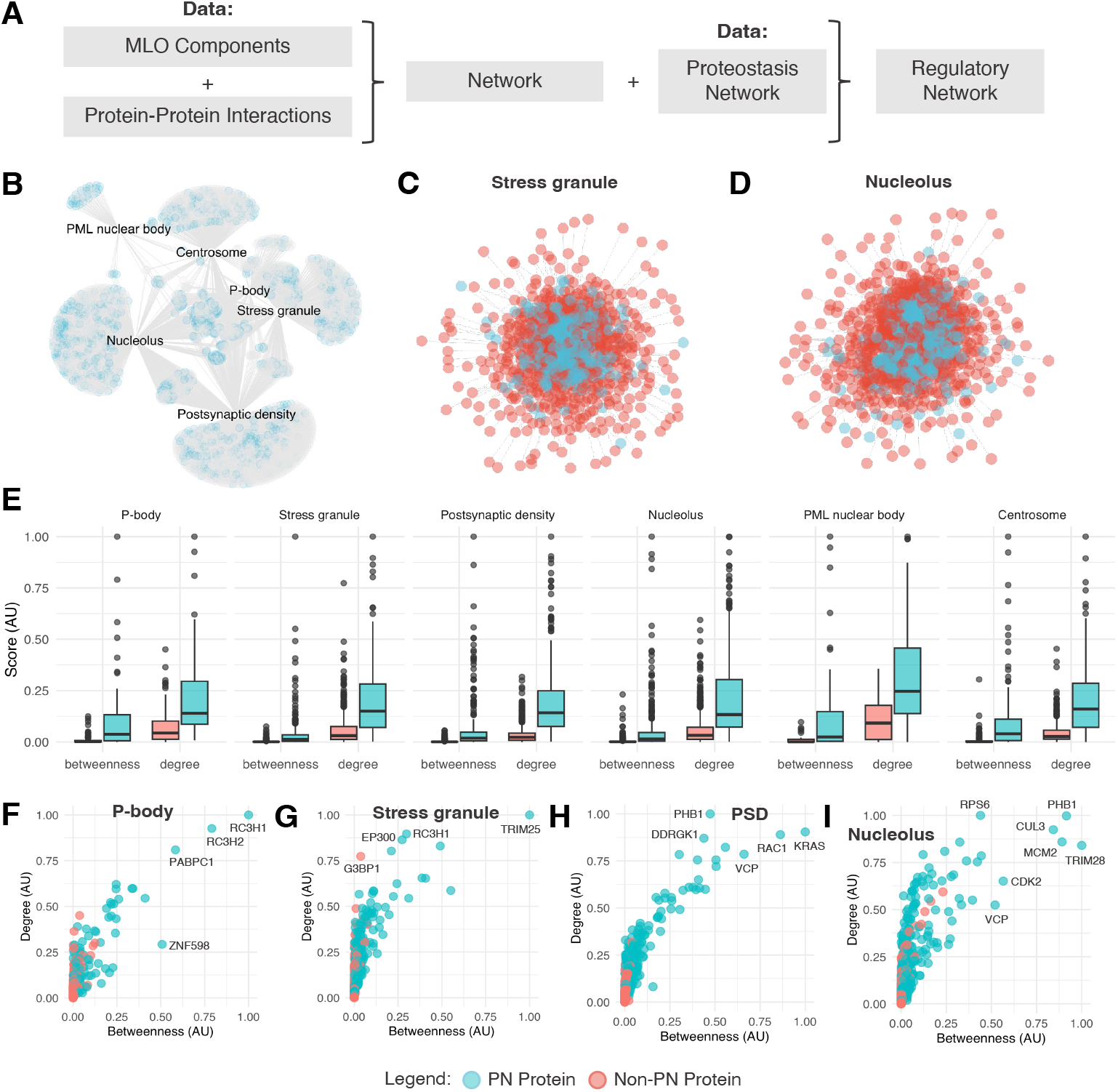
Mapping and centrality analysis of the proteostasis network (PN) of membraneless organelles (MLOs). **(A)** Flowchart illustrating the procedure used to determine the MLO-specific PNs. The protein components of a given MLO were obtained from DrLLPS ^25^, and the corresponding protein-protein interactions were extracted from BioGRID ^24^. The subset of protein-protein interactions involving PN components were then identified as the MLO-specific PN, i.e. the regulatory network of an MLO. **(B)** Overall organization of the PN that regulates MLOs (PML nuclear bodies, centrosomes, stress granules, P-bodies, nucleoli, and post-synaptic densities). Blue nodes in the protein-protein interaction network represent PN proteins and grey nodes represent non-PN proteins. Many PN proteins are involved in multiple MLOs. The full list of proteins and interactions is provided in **Supplementary Data 1. (C**,**D)** Schematic illustration of representative MLO-specific PNs for stress granules (C) and nucleoli (D). Blue nodes in the protein-protein interaction network represent PN proteins and red nodes represent non-PN proteins. In these regulatory networks, PN proteins tend to be found in the centre of the network, suggesting their essentiality within these MLO-specific PNs. **(E)** Centrality scores, including betweenness and degree, are calculated and compared for PN vs non-PN proteins within each MLO-specific PNs. PN proteins have higher degree and betweenness compared to non-PN proteins indicating their importance in network regulation. **(F-H)** PN proteins tend to be hubs and bottlenecks (hub: high degree; bottlenecks: high betweenness) of in representative MLOs - P-bodies (F), stress granules (G), post-synaptic densities (H), and nucleoli (I).

Since the data that we used are from various sources, we considered their consistency. For example, we found an overlap in the PNs of SGs and P-bodies, as can be expected due to reported similarities in their composition and function. We also analysed the unexpected observation that nucleoli and PSDs share a high number of PN components, finding that the overlapping proteins were molecular chaperones (∼22%), ribosomal proteins (∼17%), and PN proteins involved in autophagic processes (∼30%), a result consistent with previous literature reports showing that molecular chaperones are involved in the assembly of these large protein assemblies^26^. In addition, since nucleoli are the sites of ribosomal RNA (rRNA) production and assembly of ribosomal subunits, the number of nucleoli is proportional to the amount of cellular protein synthesis^27^. Furthermore, protein synthesis takes place in postsynaptic components^28^, where mRNAs and ribosomes localise in dendrites and axons to enable synaptic plasticity^29^. Also, we note that the nucleolus is a central hub for autophagic stress response^30^ and that autophagy controls synaptic transmission via postsynaptic mechanisms^31^. These observations offer a functional basis for the shared PN proteins and a mechanistic synergy between the nucleolus and synaptic compartments that shape ribosome biogenesis^32^.

### PN proteins occupy central positions in the PPI networks of MLOs

We measured and compared the roles of PN proteins against non-PN proteins within the overall PPI network of MLOs. The centrality of each PN protein was determined by two network measures (see Methods): (1) the degree, i.e. the number of connections of a protein within the network, and (2) the betweenness, i.e. the extent to which a protein lies on the shortest path between protein pairs within the network. These two measures are useful because hub proteins (high degree) and bottleneck proteins (high betweenness) were shown to play crucial topological and functional roles in maintaining the integrity and functionality of protein-protein interaction networks^33^. Our analysis reveals that PN proteins tend to have higher degrees and higher betweenness within MLO PPI networks (**Figure S2E**).

In these regulatory networks, PN proteins tend to be found in central positions in the network, as shown for representative MLO-specific PN for stress granules (**Figure 1C**) and nucleoli (**Figure 1D**). A comparison of centrality values between PN and non-PN proteins revealed that PN proteins are central in 6 of the 7 MLOs included in this study (**Figure 1E**). A twin-axis landscape of degree and betweenness is presented for representative MLOs: P-bodies (**Figure 1F**), stress granules (**Figure 1G**), post-synaptic densities (**Figure 1H**), and nucleoli (**Figure 1I**). These results highlight that: (1) hubs that are not bottlenecks tend to be structural proteins^34^, for example G3BP1 which is a key scaffold of stress granules (**Figure 1G**), and (2) hubs that are also bottlenecks tend to be part of signal transduction pathways^34^. Furthermore, proteins that are both hub and bottlenecks disrupt strongly the network upon hub removal^34^.

### Functional branches of the PN of MLOs

To study the organization of the PN of MLOs, we calculated the size of the different PN branches (PN branches) relative to the overall PN^8,9^ by dividing the number of proteins in each PN branch divided by the total number of proteins in the PN. We then calculated the size of the different PN branches relative to the PN of MLOs by dividing the number of proteins in MLOs in each PN branch by total number of proteins in the PN of MLOs. We found that the regulation of protein synthesis in the cytosol (‘Cytosolic translation’) is predominantly carried out by the PN of MLOs (**Figure 2A**). Similarly, the regulation at the subcellular level (‘Mitochondrial proteostasis’, ‘Nuclear proteostasis’ and Cytonuclear proteostasis’) is also predominantly carried out by the PN of MLOs (**Figure 2A**). By contrast, the regulation of protein degradation (‘UPS’) involves only in a limited manner the PN of MLOs (**Figure 2A**).

**Figure 2.**
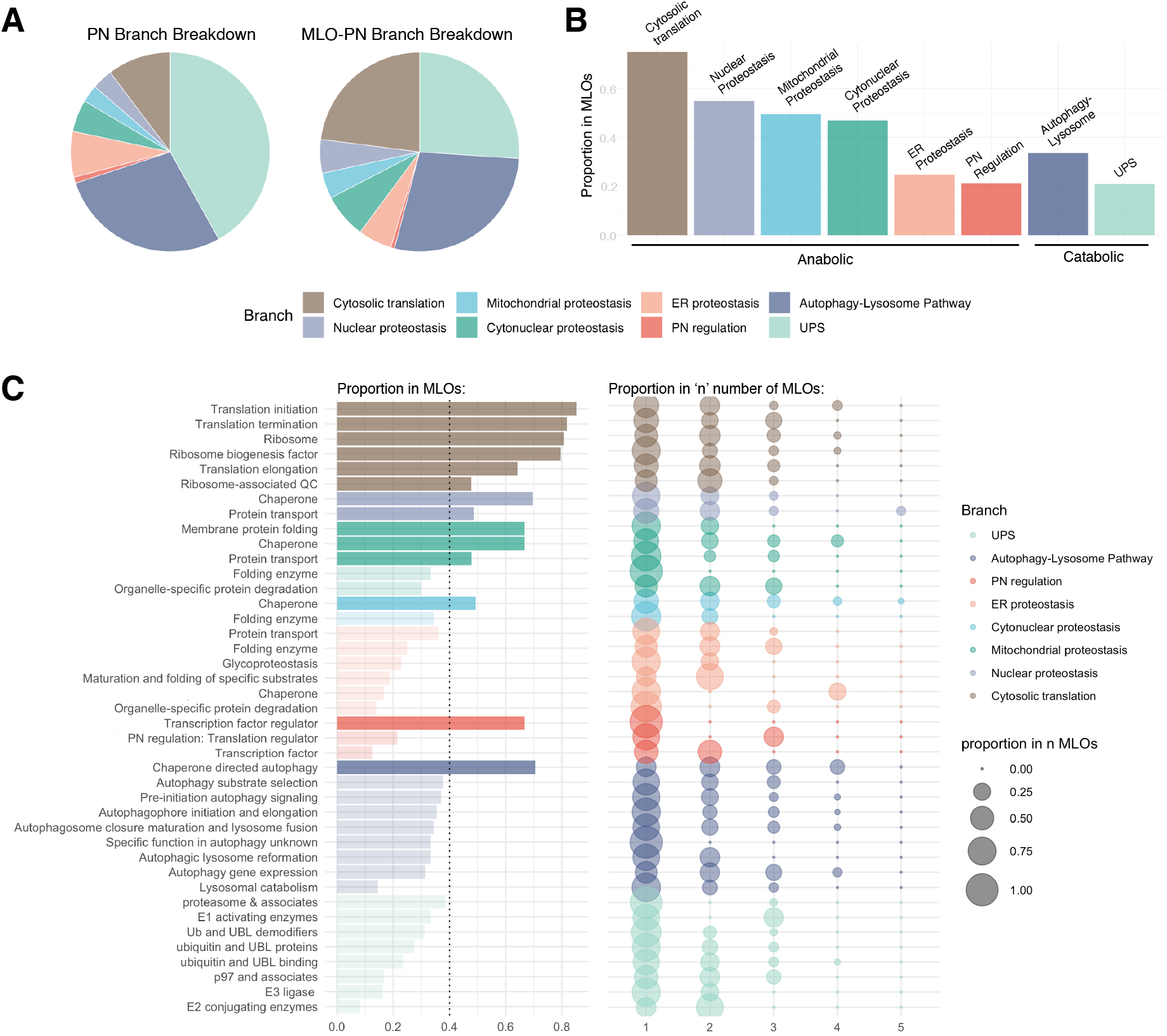
Analysis of the functional branches and functional classes of the PN of MLOs. **(A)** Comparison of the distribution of the various functional PN branches across the overall PN and the MLO-specific PN. **(B)** Proportion of each PN branch involved in MLOs. A larger proportion of anabolic PN proteins are involved in MLOs compared to catabolic PN proteins. **(C)** Proportion of each PN class involved in MLOs. A large proportion of translation-related proteins are involved in MLOs. In addition, molecular chaperones are also largely involved in MLOs. To determine the specificity of the PN classes, a bubble plot showing the proportion of proteins involved in multiple MLOs.

To support these conclusions, for each PN branch, we calculated the number of proteins in the PN of MLOs divided by number of proteins in the PN (**Figure 2B**). We found that nearly 80% of the regulation of protein synthesis in the cytosol (‘Cytosolic translation’) is associated with MLOs (**Figure 2B**). Similarly, we found that about 50% of the subcellular proteostasis (‘Mitochondrial proteostasis’, ‘Nuclear proteostasis’ and Cytonuclear proteostasis’) is associated with MLOs (**Figure 2B**). By contrast, we found that the catabolic proteostasis (‘Autophagy-Lysosome Pathway’ and ‘UPS’), which makes up of 70% of the overall PN, is not prevalently associated with MLOs (**Figure 2B**).

### Functional classes of the PN of MLOs

To continue the study of the organization of the PN of MLOs, we analysed the PN classes, which is the level below the PN branch in the taxonomy scheme of the PN^8,9^. For each PN class, we calculated the number of proteins in that class in the PN of MLOs divided by number of proteins in that class in the PN (**Figure 2C**, left). We found that several PN classes involved in protein synthesis (‘Transcription factor regulator’, ‘Translation initiation’, ‘Translation termination’, ‘Translation elongation, ‘Ribosome’, and ‘Ribosome-associated QC’) are closely associated with MLOs.

For each PN class, we then calculated the number of proteins found in one or more (n = 1, 2, 3, 4 or 5) MLOs divided by the number of proteins from that class in MLOs (**Figure 2C**, right). We found that while most PN proteins tend to be involved in only one or two MLOs, some PN proteins are involved as many as 5 MLOs. We highlight that several molecular chaperones tend to be more promiscuous than others, with HSPA2 localising in 5 MLOs (stress granules, P-bodies, nucleolus, centrosome, and PSDs), HSPA8/HSPA9 localising in 4 MLOs (stress granules, nucleolus, centrosome, and PSDs), and HSPA5 in 3 MLOs (stress granules, nucleolus, and PSDs).

### Similarity of MLO-specific PNs

Given that many PN proteins were found to localise in multiple MLOs, we measured the similarity of the 7 MLOsanalyzed in this work based on their PN components. Using the Jaccard index to measure the similarity between sets of proteins, we found that stress granules and P-bodies are most similar in terms of their MLO-specific PN proteins, followed by stress granules and nucleoli, and nucleoli and post-synaptic densities. By contrast, PML nuclear bodies share only a few components with the other 6 MLOs analyzed here (**Figure 3A**). Then, we investigated the similarity between the 3 MLOs that share most components (stress granules, nucleoli and post-synaptic densities), in terms of proteostasis function, by calculating the similarity between PN branches (**Figure 3B**). We found that nucleoli and post-synaptic densities share two main PN branches (‘cytonuclear proteostasis’ and ‘PN regulation’), that stress granules and nucleoli share one main branch (‘PN regulation’), and that the mitochondrial PN branch is shared among all 3 MLOs (**Figure 3B**). We then repeated the analysis for PN classes, showing that all 3 MLOs share ‘Autophagy-Lysosome Pathway: Chaperone-directed autophagy’ and ‘UPS: E1 activating enzymes’, but that only stress granules and post-synaptic densities share ‘UPS: E2 activating enzymes’ (**Figure 3C**).

**Figure 3.**
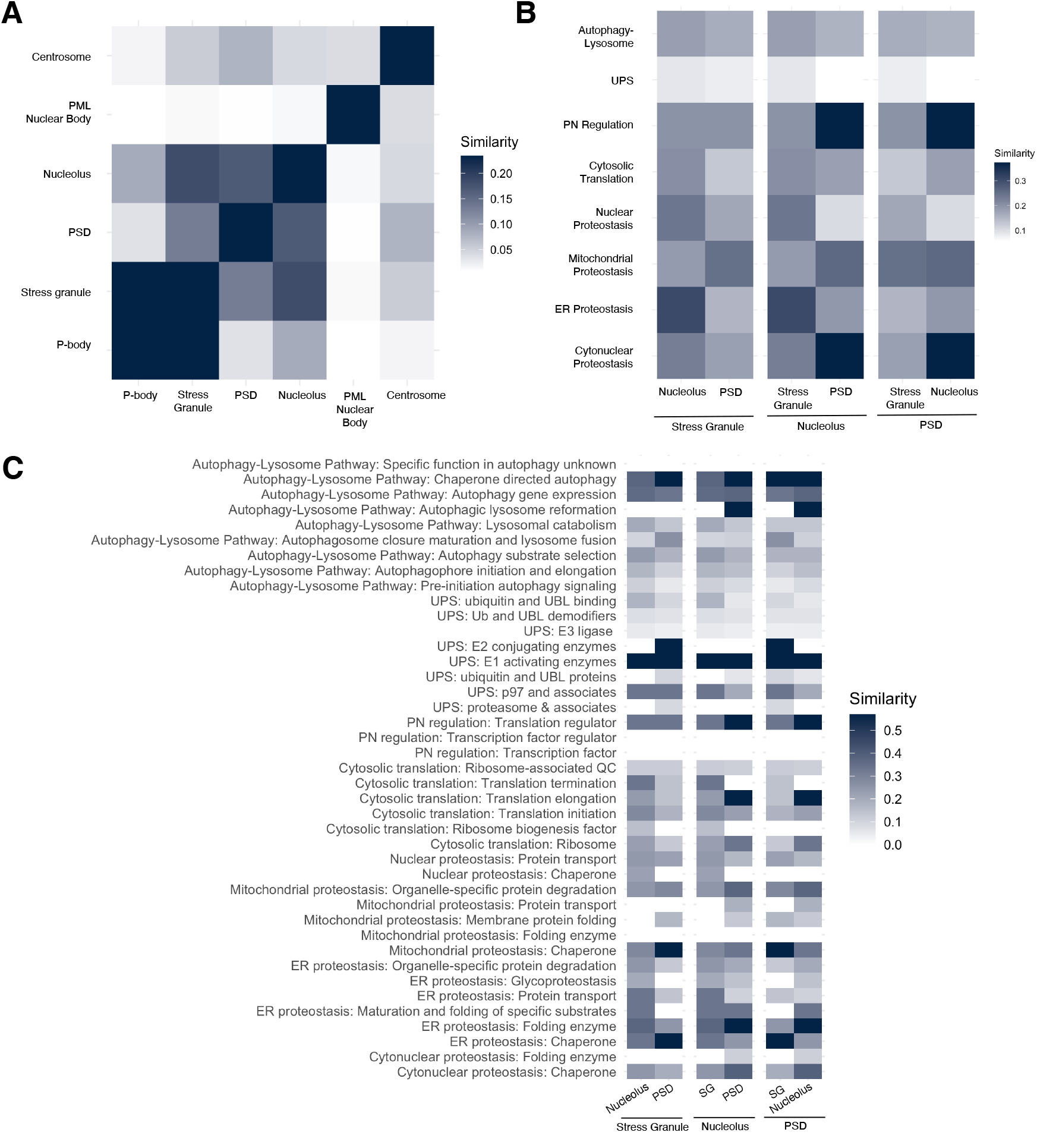
Similarity of MLOs based on their PN components. **(A)** Similarity matrix of MLOs based the number of shared MLO-specific PN components, as measured by the Jaccard index. Stress granules and P-bodies are most similar in terms of their MLO-specific PN proteins, followed by stress granules and nucleoli, and nucleoli and post-synaptic densities. By contrast, PML nuclear bodies share only a few components with the other six MLOs analyzed here. **(B)** Similarity scores between MLOs at the PN branch level. Nucleoli and post-synaptic densities share two main PN branches (‘cytonuclear proteostasis’ and ‘PN regulation’), stress granules and nucleoli share one main branch (‘PN regulation’), and the ‘mitochondrial proteostasis’ branch is shared among all three MLOs. **(C)** Similarity scores between MLOs at the PN class level. All three MLOs share ‘Autophagy-Lysosome Pathway: Chaperone-directed autophagy’ and ‘UPS: E1 activating enzymes’, but only stress granules and post-synaptic densities share ‘UPS: E2 activating enzymes’.

### HSP70s are generic while HSP70 co-chaperones are specific in the regulation of MLOs

To investigate how specificity of MLO regulation is enabled given that PN proteins co-localise to multiple MLOs and that MLOs share a large proportion of PN proteins. For this purpose, we analysed 2 hierarchial families of proteins known to exhibit client substrate specificity - molecular chaperones and E1, E2 and E3 ligases. We found that several molecular chaperones (approximately 40 out of 220) are shared across multiple MLOs (**Figure S3A,B**) while E1, E2 and E3 ligases retain the trend of substrate-specificity by being more MLO-specific, with few ligases common to multiple MLOs (**Figure S3A**). Hence, we proceed to further clarify how molecular chaperones, which were shown earlier to be closely associated with MLOs, enable substrate specificity.

To clarify how molecular chaperones enable specificity in MLO regulation, we studied the HSP70 systems^11,18^, which assist the folding and binding of client proteins. The functional properties of the HSP70 systems are critically dependent upon HSP70 co-chaperones to bind and execute their function on substrate proteins. We hence hypothesized that HSP70 chaperones may have broad regulatory functions on multiple MLOs while co-chaperones may have MLO-specific regulatory capabilities. This assumption is based on earlier reports that HSP70 inhibition impacts MLO dynamics^19,35^. To test this hypothesis, we first mapped out the localization of HSP70 chaperones and their co-chaperones. Our localization map revealed that HSP70 proteins are indeed promiscuous to multiple MLOs, while HSP70 co-chaperones tend to localize to different types of MLOs (**Figure 4A**).

**Figure 4.**
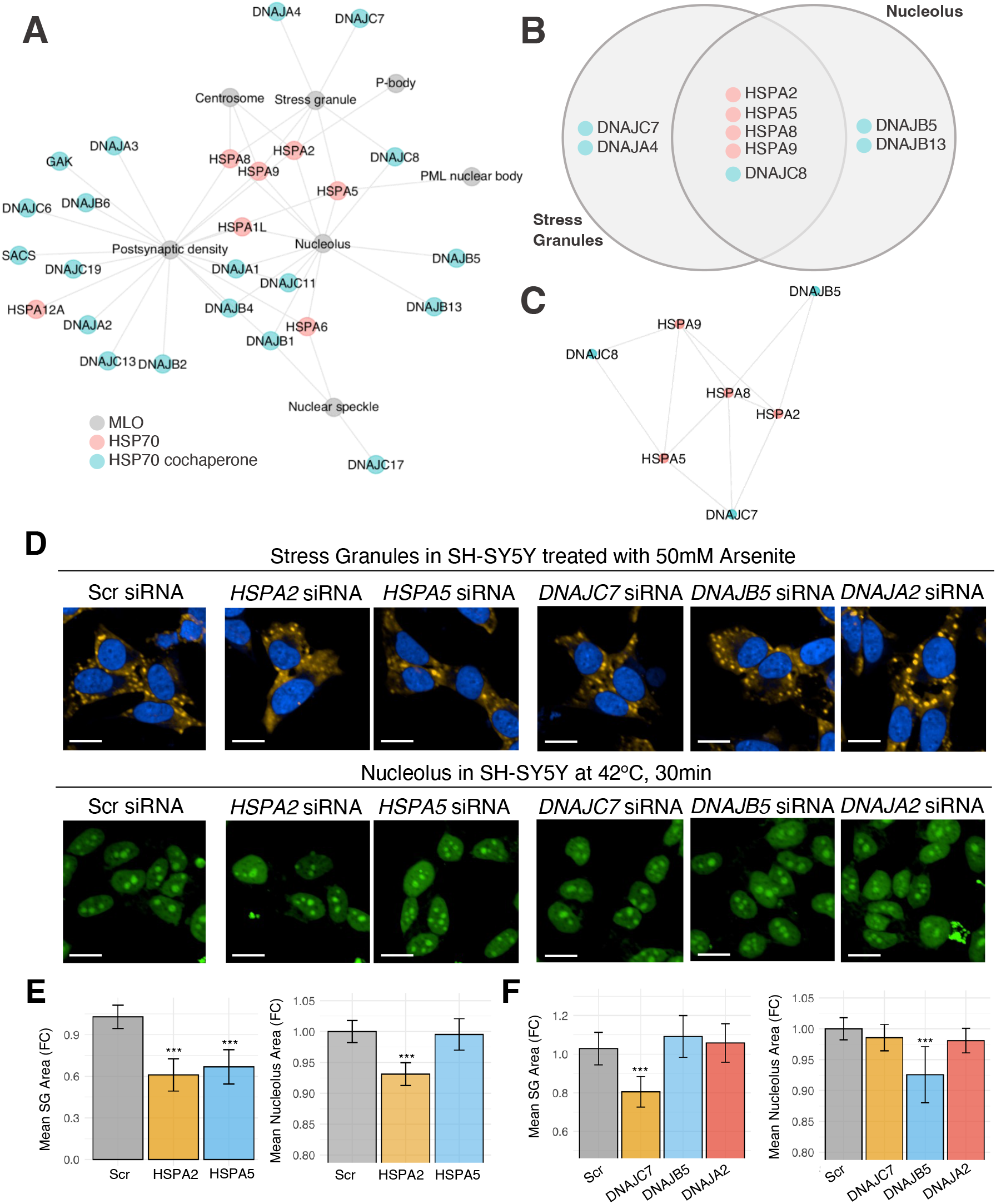
HSP70 proteins regulate multiple MLOs while their co-chaperones are specific to individual MLOs. **(A)** HSP70 chaperones (pink) tend to be present in multiple MLOs, while HSP70 co-chaperones (teal) tend to be more specific in their MLO localisation. **(B)** HSPA2, HSPA5, HSPA8, and HSPA9 are HSP70 chaperones present in both stress granules and nucleoli. In contrast, while DNAJC8 is common to both MLOs, DNAJC7 and DNAJA4 localize in stress granules while DNAJB5 and DNAJB13 are localising in nucleoli. **(C)** Interaction network of HSP70 proteins common to stress granules and nucleoli, and selected co-chaperones. **(D)** Representative images of stress granules formed when SH-SY5Y cells are treated with 0.5 mM of sodium arsenite for 30 min, and nucleoli observed upon heat stress at 42 °C for 30 min. **(E)** Upon treatment with *HSPA2* siRNA, we observed a decrease in the average stress granule and nucleoli areas. Similarly, upon treatment with *HSPA5* siRNA, average stress granule area decreased significantly. **(F)** Average stress granule area only changes (decreases) upon treatment with *DNAJC7* siRNA and not *DNAJB5* or *DNAJA2* siRNA. Average nucleoli area only changes (decreases) upon treatment with *DNAJB5* siRNA and not *DNAJC7* or *DNAJA2* siRNA. N = 3 biological replicates of n = 6 technical replicates (for stress granules), and n = 3 technical replicates (for nucleoli).

Analyzing stress granules and nucleoli as case studies, we found that they share four HSP70 chaperones (HSPA2, HSPA5, HSPA8, and HSPA9), but only share one HSP70 co-chaperone (DNAJC8, **Figure 4B**). Interactions between the shared HSP70 chaperones and three types of HSP70 co-chaperones (SG-specific, nucleoli-specific, and shared) are presented (**Figure 4C**).

For experimental validation, we selected the following proteins: (i) HSPA2 and HSPA5, which are HSP70 chaperones common to both stress granules and nucleoli, (ii) DNAJC7, which is a SG-specific HSP70 co-chaperone, (iii) DNAJB5, which is a nucleolus-specific HSP70 co-chaperone, and (iv) DNAJA2, a HSP70 co-chaperone taken as negative control as it localizes to post-synaptic densities within the MLO-specific PN. Within our experimental assay, we individually knocked down HSPA2, HSPA5, DNAJC7, DNAJB5, and DNAJA2 in SH-SY5Y cells using the corresponding siRNA for 72 h. We then studied the changes in stress granules and nucleoli formation through triggering changes in their formation by treating the cells with 50 μM sodium arsenite for 30 min or heating the cells at 42°C for 30 min. Based on our hypothesis, we expected to see changes in both stress granules and nucleoli upon HSP70 protein knockdown, but MLO-specific changes upon knockdown of HSP70 co-chaperones, with the exception of the negative control DNAJA2.

Our experimental results confirmed our hypothesis that HSP70 chaperones are promiscuous while HSP70 co-chaperones are specific to MLO regulation. Based on our quantification, treatment with *HSPA2* siRNA decreased the formation of stress granules and nucleoli upon 50 μM sodium arsenite and heat shock treatment (**Figure 4D**), confirming the promiscuous activity of HSP70 chaperones across MLOs. Similarly, treatment with *HSPA5* siRNA also resulted in decreased total area of stress granules formed within each cell (**Figure 4D**,**E**).

In contrast, treatment with siRNAs targeting HSP70 co-chaperones resulted in higher specificity at targeting MLO formation. Treatment with *DNAJC7* siRNA resulted in a lowering of the total area of stress granules formed but not the total area of nucleoli formed. In addition, treatment with *DNAJB5* siRNA resulted in a lower total area of nucleoli formed but not the total area of the stress granules formed. Finally, treatment with our negative control *DNAJA2* siRNA did not affect the total area of stress granules or nucleoli formed within SH-SY5Y cells (**Figure 4D,F**).

## Discussion and Conclusions

In this work, we sought to understand the organization of the proteostasis system of MLOs by asking how different MLOs achieve functional specificity in protein quality control despite an overlap in their components. To address this question, we mapped the PN of membraneless organelles and analyzed its organization. We thus identified the major components of this PN and measured the similarity of different MLOs in terms of their proteostasis regulation, finding several shared mechanisms.

Next, to understand how the proteostasis system achieves regulation of MLOs despite these similarities, we explored how the PN components are capable of specifically modulating different membraneless organelles. We showed how HSP70 chaperones are promiscuous in their regulatory activity across MLOs, while the co-chaperones of HSP70 enable specificity in targeting MLOs.

Our results revealed the extent to which one of the key functions of molecular chaperones is to regulate the conformational behaviour of proteins not just during folding and binding, as well as misfolding and aggregation, but also within MLOs.

The comprehensive mapping that we provided of the proteostasis system of MLOs will enable future investigations of several important aspects of the behaviour of MLOs:

*Formation mechanisms*. Increasing evidence suggests that MLOs are formed by a phase separation mechanism^3,4^. It is still unclear, however, what are the exact molecular and biophysical determinants that drive the phase separation process leading to the formation of distinct MLOs.

*Response to environmental changes*. It will also be important to achieve a better understanding of how the assembly and disassembly of MLOs are dynamically regulated during various cellular conditions, such as stress or changes in metabolism^12^. Understanding the triggers and mechanisms underlying these transitions would provide insights into the adaptability of the PN.

*Cross-talk and coordination*. There is a need to reveal the nature of the cross-talk between different MLOs and the mechanisms by which they coordinate their roles within the PN.

*Role of RNA in the self-assembly and function:* RNA is a critical component in many MLOs, as RNA molecules can influence the dynamics, composition, and function of MLOs^36,37^. Detailed investigations of how RNA interacts with proteins within the PN to regulate the properties and functions of MLOs will provide a more comprehensive view of their regulation.

*Pathological aggregation*. It remains to be clarified how the dysregulation of the self-assembly process of MLOs leads to the pathological aggregation of proteins associated with neurodegenerative diseases, such as Alzheimer’s disease and amyotrophic lateral sclerosis (ALS)^4,6,7^, and how the PN may help prevent this phenomenon^20,38^.

We anticipate that further studies based on the mapping of the proteostasis system of MLOs will uncover more detailed mechanisms of the proteostasis regulation of these cellular structures, thus offering new avenues for target identification for therapeutic interventions for diseases associated with MLOs.

## Methods

### Proteostasis network (PN) and protein-protein interaction (PPI) data

A list of annotated PN components^8,9^ was obtained from the Proteostasis Consortium (https://www.proteostasisconsortium.com/). *Homo sapiens* MLO components with annotated scaffolds were downloaded DrLLPS^25^. *Homo sapiens* PPI data were downloaded from BioGRID^24^ (accession date: Nov2023). Paired interactions from BioGRID were filtered for protein pairs that physically coexist (here-after referred to as ‘physical-PPI pairs’) using tags: “MI:0407 (direct interaction)”, “MI:0915 (physical interaction)”, “MI:0914 (association)”, and “MI:0403 (colocalization)”.

### Selection of key MLOs for study

The number of proteins characterised were computed for each MLO documented in the DrLLPS^25^ database. 7 well-characterized MLOs containing more than 50 proteins were selected for this study. The 7 key MLOs selected are: nucleoli, stress granules, P-bodies, centrosomes, nuclear speckles, PML nuclear bodies (PML-NB), and post-synaptic densities (PSD).

### Identifying MLO-specific PN-networks

For each MLO include in the study, we set up a MLO-specific PN network by integrating DrLLPS^25^ and the filtered BioGrid data. An MLO-specific PPI edgelist was generated by filtering the BioGrid physical-PPI pairs for interaction pairs that involved two proteins reported in the DrLLPS dataset to localise within the MLO. The edgelist was then used to construct a non-directional PPI network specific to each of the key MLOs. The edgelist for the PPI networks is available in **Supplementary Data 1**.

### MLO PN proteins vs non-PN proteins

Proteins within the PN network of each MLO were divided into 2 groups: PN proteins and non-PN proteins. PN proteins within the protein network of an MLO are PN components that are part of the PN network of the MLO. Non-PN proteins are the complementary set of proteins to PN proteins within the PN network of an MLO.

### Identification of hub proteins and bottleneck proteins in MLOs

To identify essential proteins in the MLO-specific PNs using quantifiable metrics, we determined the relative ‘hubness’ and ‘bottleneckness’ of individual PN components within the MLO-specific PNs. Hub proteins are proteins with a high degree within the network and protein bottlenecks are proteins with high betweenness^34^. The degree centrality (degree) of a node in a network refers to the number of its edges^39^. Hence, in our analysis of MLO-specific PNs, the degree of a protein refers to the number of neighbouring proteins it interacts with. The betweenness centrality (betweenness) of a node quantifies the number of shortest paths passing through the node^36,40^. Nodes with high betweenness are critical points within a network determining the amount of information flow and are defined as bottleneck proteins^34^. Bottleneck proteins are essential proteins, as failure of bottleneck proteins result in the collapse of the biological network. Hub-bottlenecks hence refer to proteins with both high degree and betweenness^34^. Centrality scores are available in **Supplementary Data 2**.

### Similarity of membraneless organelles based on their PN protein components

The Jaccard index was used as a measure of similarity of MLOs based on their constitutive PN components. The Jaccard index is defined here as the number of shared PN components divided by the total number of PN components within both MLOs.

### Cell culture experiment

Human neuroblastoma cells (SH-SY5Y) were cultured in DMEM/F-12 GlutaMAX™ supplemented with 10% heat-inactivated fetal bovine serum (hiFBS) and maintained at 37 °C, 5% CO2, 95% relative humidity. For each experiment, SH-SY5Y cells were plated in PerkinElmer Cell Carrier Ultra plates at a density of 7.5k-10k cells/well. The cells were incubated for 24 h to allow adequate cell attachment before siRNA feeding. During siRNA feeding, full culture medium was replaced with fresh DMEM/F-12 GlutaMAX™ with 1% hiFBS and fed with a final concentration of 10 nM siRNA. Lipofectamine RNAiMAX was used as the transfection reagent. The cells were incubated with Thermo Fisher silencer select siRNAs (HSPA2: #s20124 & #s20122, HSPA5: #s6979 & #s6980, DNAJC7: #s270217, DNAJB5: #s24554 & #s24556, and DNAJA2: #s6973 & #s7975) for 72 h before treatment with sodium arsenite or heat shock. siRNA knockdown efficiency for all genes were confirmed via PCR (**Figure S4**). For sodium arsenite treatment, cells were supplied with fresh DMEM/F-12 GlutaMAX™, 1% hiFBS with 50 μM sodium arsenite and left to incubate for 30 min at 37 °C, 5% CO2, 95% relative humidity before fixing. For heat shock treatment, cells were incubated on a pre-heated hot plate at 42 °C for 30 min before fixing. The SH-SY5Y cells were fixed using 4% paraformaldehyde (PFA) dissolved in D-PBS (+/+) and stored at 4 °C for less than a week before staining.

### RNA extraction and reverse transcription

Total RNA was extracted using PureLink™ RNA Mini Kit (#12183018, Invitrogen) according to the manufacturer’s protocol. 700ng to 1μg of RNA was reverse transcribed into cDNA in a 20 μL volume using the High-Capacity cDNA Reverse Transcription Kit (#4368814, Applied Biosystems) according to the manufacturer’s protocol.

### Quantitative PCR

The qRT-PCR (HSPA5, DNAJC7, DNAJB5, and DNAJA2) was carried out in a volume of 20μl, which contained 1μl cDNA, 10μl TaqMan™ Gene Expression Master Mix (#4369016, Applied Biosystems), and 1x of each Taqman probes (ThermoFisher). The following reaction conditions were employed according to the manufacture’s instruction: 50 °C for 2 min, 95 °C for 10 min, then 40 cycles of 95 °C for 15 sec and 60 °C for 1min. The relative level of mRNA was evaluated with the ΔΔCt method, using GAPDH as endogenous control.

A preamplification reaction was carried out for HSPA2 in a volume of 20 μl, which contained 4 μl cDNA, 10 μl TaqMan™ Gene Expression Master Mix (#4369016, Applied Biosystems), and 0.05x of each Taqman probes (ThermoFisher). The following reaction conditions were employed according to the manufacture’s instruction: 95 °C for 10 min, then 10 cycles of 95 °C for 15 sec and 60 °C for 4 min. The qRT-PCR (HSPA2) was then carried out in a volume of 20 μl, which contained 5 μl preamplification product, 10 μl TaqMan™ Gene Expression Master Mix (#4369016, Applied Biosystems), and 1x of each TaqMan probes (ThermoFisher). The following reaction conditions were employed according to the manufacture’s instruction: 50 °C for 2 min, 95 °C for 10 min, then 40 cycles of 95 °C for 15 sec and 60 °C for 1 min. The relative level of mRNA was evaluated with the ΔΔCt method, using ACTB as an endogenous control.

### Immunocytochemistry

Cells were permeabilized with 0.1% Triton™ X-100 (#85111, Thermo Scientific) for 20-30 min at room temperature then rinsed three times with D-PBS (+/+). Following which, the cells were incubated for at least 1 h at room temperature in 1.5% BSA blocking buffer. After blocking, cells were incubated with the appropriate dilution of the desired primary antibodies overnight at 4 °C. To stain for stress granules, G3BP1 primary antibody G3BP1 (#ab109012, Abcam) at 1 μg/mL, washed three times with D-PBS (+/+), and incubated with secondary antibody AlexaFluor555 anti-rabbit (#A21428, Thermo Fisher) diluted in blocking solution for 1 h at room temperature. To stain for nucleoli, the Nucleolus Bright Green dye (#NBP1-86651, Novus Biologicals) was diluted in DPBS at 1:1000. Post-staining, the cells were rinsed three more times with D-PBS (+/+) and stained with Hoechst 33342 (#H3570, Thermo Fisher) before imaging. All 96-well plates generated were imaged using the Opera Phenix High-Content Confocal microscope at a 40X magnification. Images of 16 to 20 unique frames per well (technical replicate, n=3) for each biological replicate (N=3) were acquired for quantification. Image analysis and quantifications were performed with the Harmony High-Content Imaging and Analysis Software (Perkin Elmer). G3BP1 fluorescence was used to segment and quantify stress granule area, while Nucleolus Bright Green dye staining for nucleoli RNA was used to segment and quantify nucleoli area.

## Acknowledgements

This study was funded in part by UKRI (10059436 and 10061100)

## SUPPLEMENTARY INFORMATION

**Supplementary Data 1. PN of MLOs**. The PN of MLOs is made up of over 220,000 pairwise interactions. Each interaction involves one protein in the PN (column A) and one protein reported as present in one of 7 well-characterized MLOs (PML nuclear bodies, centrosomes, stress granules, P-bodies, nucleoli, and post-synaptic densities) (column C).

**Supplementary Data 2**. Lists of the centrality scores (degree and betweenness) for all proteins in their MLO regulatory networks, for each MLO.

**Figure S1.**
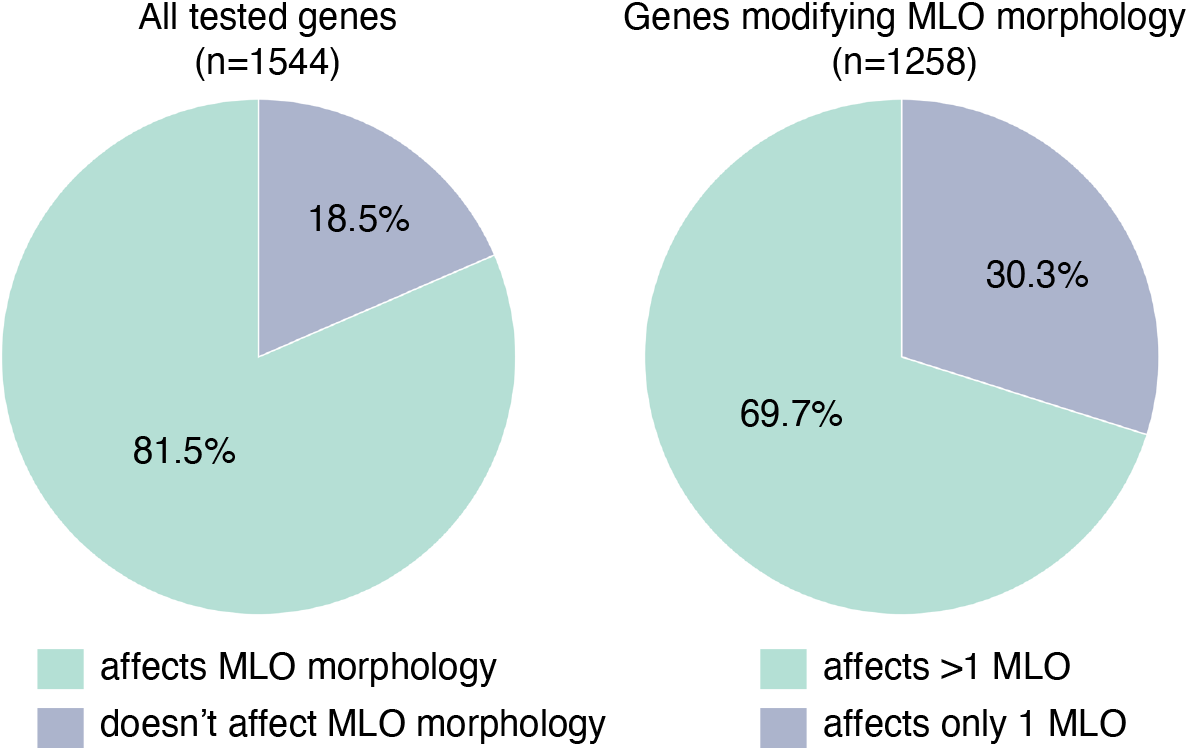
Specificity of genes previously found to affect the morphology of MLOs. In a large-scale study, a large majority (over 80%) of the of 1544 genes tested via siRNA knockdown were found to cause changes in the morphology MLOs^1^. In addition, about 70% of these genes affected the morphology of more than MLO, suggesting low specificity in the regulation of MLOs.

**Figure S2.**
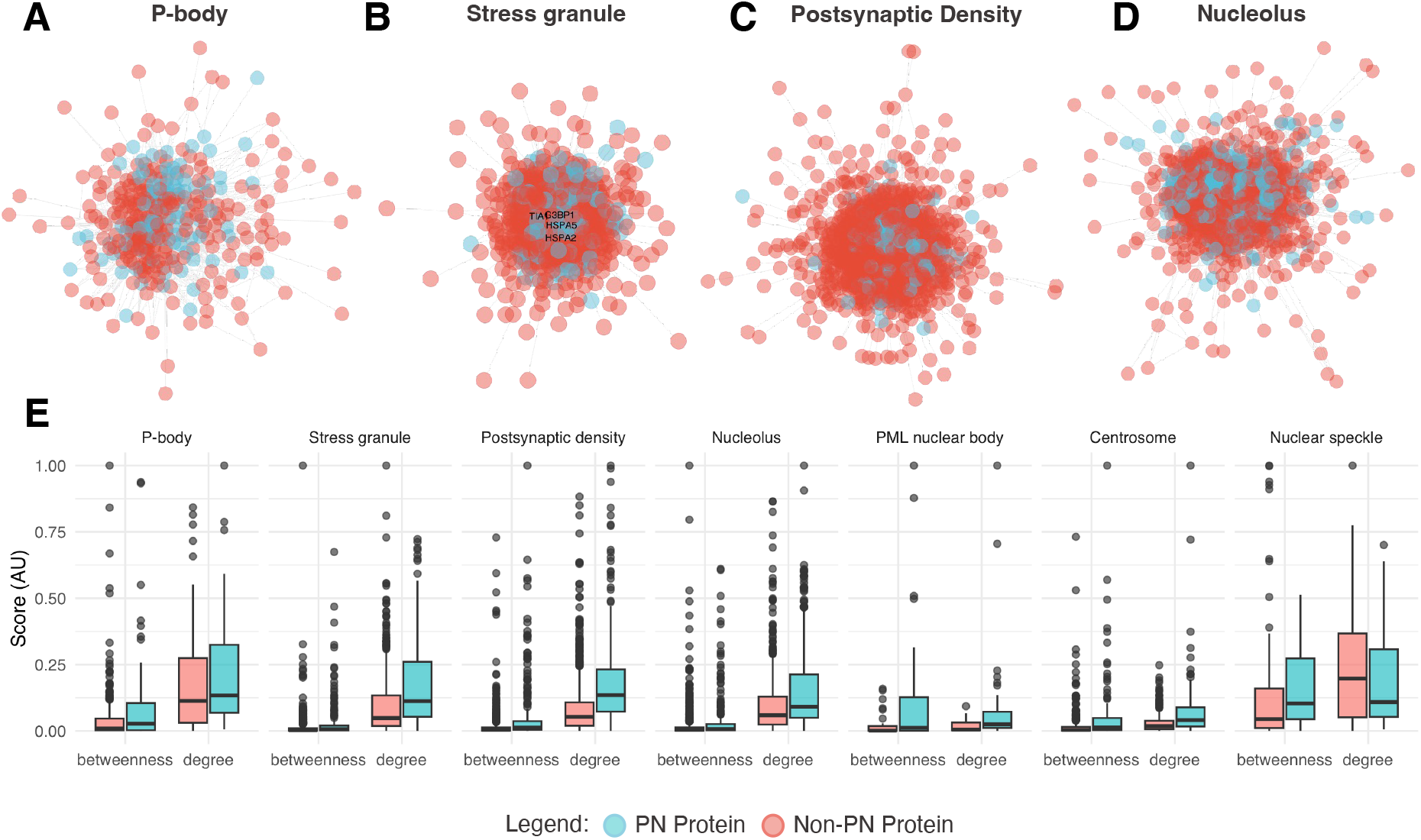
Protein-protein interaction network of the 7 MLOs analyzed in this study and centrality scores of PN proteins. **(A-D)** Overall protein-protein interaction networks for P-bodies **(A)**, stress granules **(B)**, post-synaptic densities **(C)**, and nucleoli **(D)**. Blue nodes in the protein-protein interaction network represent PN proteins and red nodes represent non-PN proteins. **(E)** Two centrality scores - betweenness and degree - are calculated and compared for PN vs non-PN proteins within each MLO PPI network. PN proteins are more central on average, suggesting regulatory importance, within P-bodies, stress granules, post-synaptic densities, and nucleoli.

**Figure S3.**
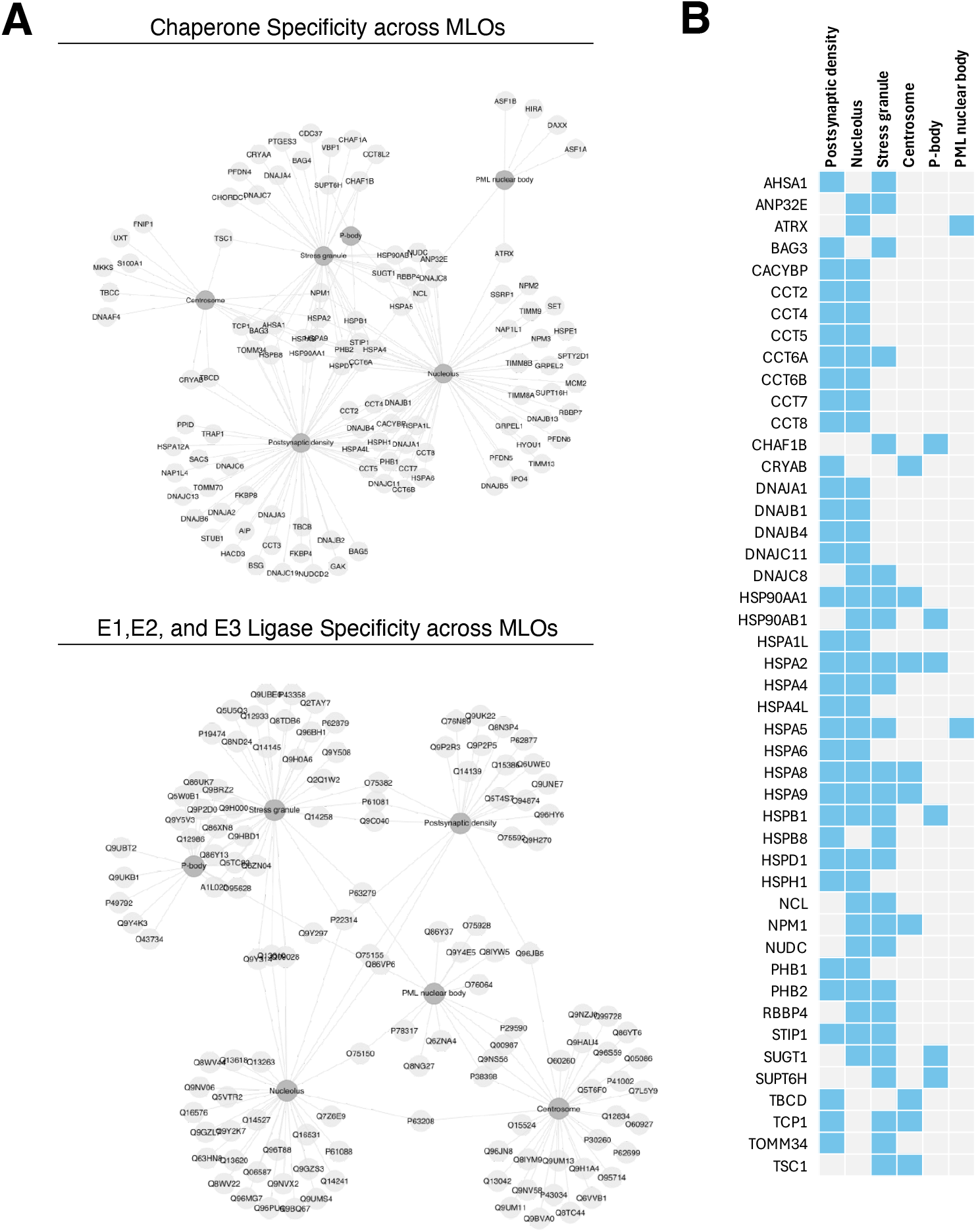
Molecular chaperones tend to be MLO promiscuous, while UPS E1, E2, and E3 ligases tend to be MLO specific. **(A)** Localisation patterns of molecular chaperones and E1, E2, and E3 ligases across MLOs. A large number of molecular chaperones (approximately 40 out of 220) are found to be shared across multiple MLOs. In contrast, E1, E2, and E3 ligases tend to be more specific to MLOs with few ligases common to multiple MLOs. **(B)** Localisation of promiscuous chaperones across MLOs. We note that in addition to the HSP70 family of proteins and their co-chaperones, several members of the small HSP family such as HSPB1 and HSPB8 appear to be promiscuous across multiple MLOs. Previous work has reported the role of HSPB8 in maintaining the dynamic of stress granules^2,3^, as well as the role of HSPB1 in both regulating the dynamics of stress granule and TDP-43 containing condensates^2,4,5^. In this work we focus on the hierarchial protein families such as HSP70 and their co-chaperones, and E1/E2/E3 ligases known to have a funneling effect on substrate specificity at lower levels.

**Figure S4.**
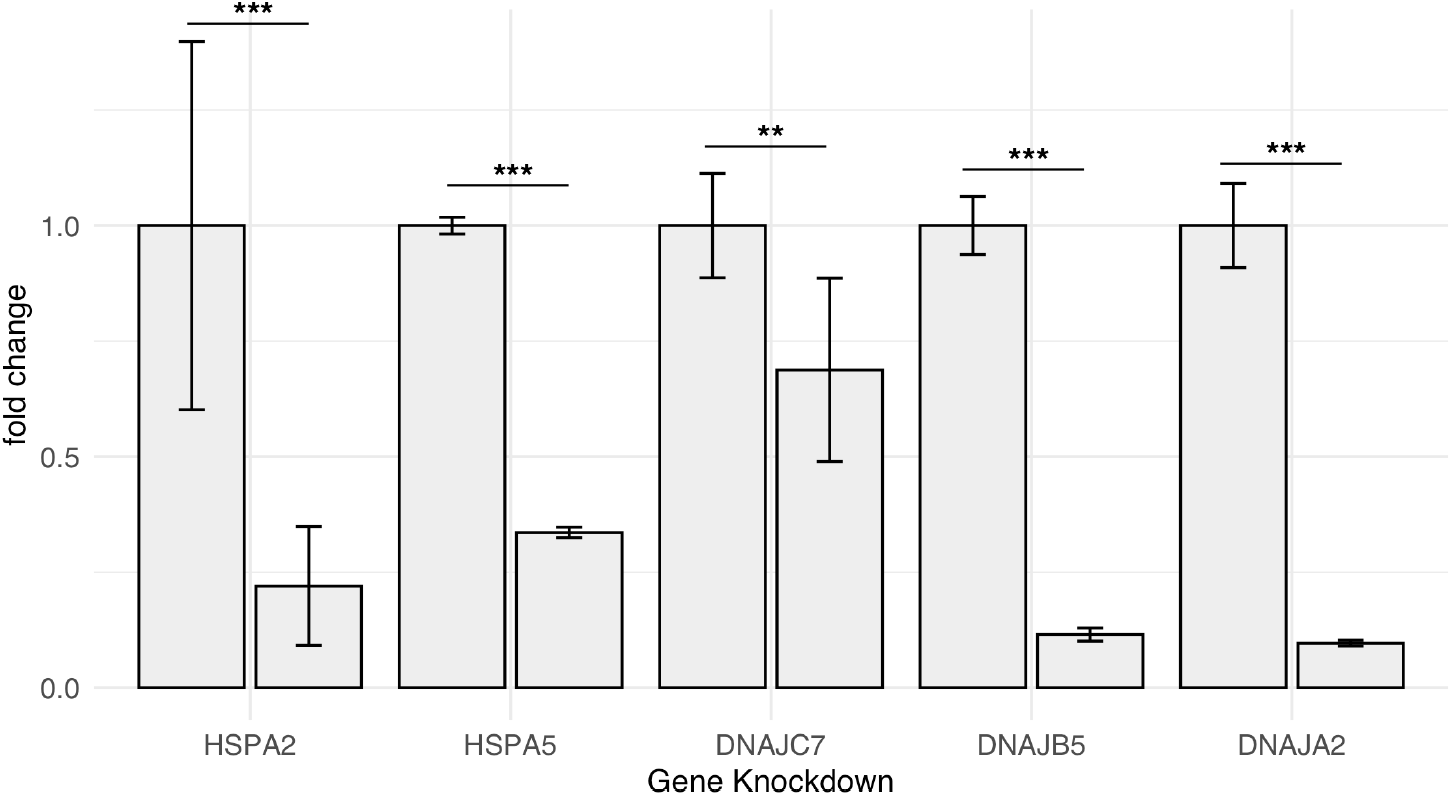
Validation of siRNA knockdown efficiency. PCR was carried out to validate the knockdown efficiency of all siRNAs used. The statistical significance was determined using the ANOVA test. (***) p-value < 0.01, and (**) p-value < 0.05.

